# Modelling genetic risk of β-cell dysfunction in human induced pluripotent stem cells from patients carrying the *MTNR1B* risk variant

**DOI:** 10.1101/2024.07.19.603674

**Authors:** T. Singh, S. Kalamajski, J.P.M.C.M. Cunha, S. Hladkou, F. Roberts, S. Gheibi, A. Soltanian, K. Yektay Farahmand, O. Ekström, A. Mamidi, P.W. Franks, A. Rosengren, H. Semb, H. Mulder, M. Fex

## Abstract

Disruptions in circadian rhythm, partly controlled by the hormone melatonin, increase the risk of type 2 diabetes (T2D). Accordingly, a variant of the gene encoding the melatonin receptor 1B (*MTNR1B*) is robustly associated with increased risk of T2D. This single nucleotide polymorphism (SNP; rs10830963; G-allele) is an expression quantitative trait locus (eQTL) in human pancreatic islets, conferring increased expression of *MTNR1B*, which is thought to perturb pancreatic β-cell function. To understand this pathogenic mechanism in detail, we utilized human induced pluripotent stem cells (hiPSC), derived from individuals with T2D carrying the *MTNR1B* G-allele. Patient-derived fibroblasts were reprogrammed to hiPSC and single-base genome editing by CRISPR/Cas9 was employed to create isogenic lines of either the C/C or G/G genotypes (non-risk and risk, respectively). In addition, the human embryonic stem cell (hESC) line (HUES4) was subjected to genome editing to create isogenic lines of either the C/C or G/G genotypes. hiPSC and hESC were differentiated into β-cells, using a 50-day 2D protocol. Single-base genome editing generated cells with the desired genotype at a success rate of >90%. Expression of stage-specific markers confirmed differentiation of both hiPSC and hESC into β-cells. *MTNR1B* mRNA levels were consistently low in differentiated β-cells, precluding quantitative analysis of gene expression. However, Western blot analysis showed higher levels of MTNR1B in differentiated β-cells carrying the risk allele, consistent with rs10830963 (G-allele) being an eQTL in β-cells. Insulin secretion in response to glucose and IBMX was similar between the genotypes, whereas addition of melatonin reduced secretion in G-allele carriers. We conclude that the stem cell-derived β-cells are not sufficiently mature to allow determination of eQTL status at the mRNA level. However, we did observe increased MTNR1B protein and increased sensitivity of β-cells from risk allele carriers (G-allele) to melatonin with regard to insulin secretion, thus supporting a functional role for the rs10830963 SNP in β-cell dysfunction.

## Introduction

Pancreatic β-cells reside in small clusters of endocrine cells, termed the islets of Langerhans, which are dispersed throughout the exocrine parenchyma. The β-cells secrete insulin, a hormone with glucose lowering effects, in response to nutrients absorbed after a meal. This maintains blood glucose levels within a narrow physiological range.

Type 2 diabetes (T2D) is a metabolic disease whose prevalence continues to increase globally and represents a major cause of morbidity and mortality. β-cell function (i.e., the capacity to secrete insulin) plays a crucial role in the development of T2D, as failure to compensate for an increased demand for insulin owing to insulin resistance, is central to the pathogenesis of the disease. Although both insulin resistance and β-cell dysfunction are hallmarks of T2D, impaired insulin secretion is the essential culprit as insulin resistance alone does not result in T2D [1–3]. This notion has been further corroborated by genome-wide association studies (GWAS) of T2D and related traits, which have identified more than 1000 robust genetic association signals [4] that mainly map to genes implicated in β-cell function [5]. One such genetic signal maps to the *MTNR1B* locus; rs10830963, a single nucleotide polymorphism (SNP) located in the intron of the *MTNR1B* gene. It is one of the most robustly replicated risk variants for hyperglycemia and diminished insulin secretion related to T2D development. Carriers of the G-risk allele exhibit decreased capacity to release insulin, increased fasting plasma glucose and a greater risk of developing T2D (or gestational diabetes) as compared to carriers of the non-risk allele C-allele [2, 6–9]. In addition, rs10830963 is an expression quantitative trait locus (eQTL) conferring increased expression of *MTNR1B* mRNA in islets from risk allele carriers [2]. Indeed, the correlation between disruptions in circadian rhythm, partly regulated by the melatonin, appears to increase the risk of type 2 diabetes (T2D) [10].

Despite the strong genetic association of the rs10830963 SNP with T2D and related traits, the precise molecular mechanisms underpinning the role of melatonin signalling in the pathogenesis of T2D have not yet been determined. To this end, we have shown that rs10830963 is an eQTL, conferring increased expression of *MTNR1B* mRNA in human islets of Langerhans [2]. Studies in clonal β-cells (INS-1 832/13) showed that melatonin reduces both insulin secretion and levels of intracellular cAMP, whereas *Mt2* (*Mt2* being the equivalent of the human *MTNR1B* gene) knockout mice display an increase in insulin secretion from isolated islets due to loss of *Mt2* signalling [6]. Indeed, the MTNR1B receptor signals via inhibitory G-proteins (G_i_), when activated by melatonin, thus abrogating insulin secretion by reducing cAMP. In addition, daily administration of melatonin accentuates the repression of insulin secretion in *MTNR1B* G/G risk allele carriers [6]. Bioinformatics and functional analyses support a pathogenic role of rs10830963, where the risk SNP introduces a putative binding site for the transcription factor NEUROD1, which potentially drives the increased expression of *MTNR1B* in β-cells [11]. While these observations collectively support a pathogenic role of melatonin signaling in T2D, a causal relationship between rs10830963 (G/G versus C/C at this position in the DNA) is yet to be established.

Here, we describe a stem cell-based approach to delineate the role of the *MTNR1B* rs10830963 risk allele in β-cell function. Homozygous individuals carrying the G-risk allele were identified and induced pluripotent stem cells (hiPSC) were generated from fibroblasts in skin biopsies. Isogenic, homozygous non-risk C-allele hiPSC, were created by single-base genome editing. Additionally, human embryonic stem cells (hESC) were genome-edited to generate cells homozygous for risk (G) and non-risk (C) alleles. Subsequently, cells carrying risk- and non-risk alleles were transcriptionally, morphologically, and functionally compared to gain information about a potential causal relationship between rs10830963 in β-cells and T2D.

## Material and Methods

### Ethical statement

All participants provided informed consent for donating a skin biopsy. The study was approved by the Swedish Ethical Review Authority.

### Preparation of human dermal fibroblasts from adult skin biopsies

Patients were recruited from “Detailed assessment of T2D” (DIACT), a sub-group of ANDIS (All new diabetics in Scania, https://www.andis.lu.se/startsida). A 4.0 mm skin punch biopsy was collected, cut into 0.5 to 1 mm pieces, and placed in a 6-well plate containing medium (DMEM, 10% FBS, Pen/Strep, Sigma Aldrich, D6429). A cover slip was placed over the biopsy pieces to fix them to the bottom of the plate. Following incubation for 7 - 10 days at 37 °C, 5% CO_2_, a dense outgrowth of fibroblasts appeared which adhered to the cover slip. Fibroblasts from the cover slip were collected, using trypsin, and further expanded in T-25 tissue culture flasks as described [12]. Cells were stored in liquid nitrogen or were directly subjected to reprogramming into iPSC.

### Reprogramming of dermal fibroblasts to iPSC

Reprogramming into hiPSCs was performed in a complete xeno-free culture environment, using non-integrating, non-modified RNAs from a highly efficient and robust Stemgent StemRNA 3rd Gen Reprogramming kit (REPROCELL, Cat. No. 00-0076). This RNA-based method combines a cocktail of synthetic, non-modified reprogramming (OCT4, SOX2, KLF4, cMYC, NANOG, and LIN28) and immune evasion mRNAs (E3, K3, B18) with reprogramming-enhancing mature, double-stranded microRNAs from the 302/367 cluster. Use of live staining with Stain Alive TRA-1-60 antibody enabled verification and selection of pluripotent clones. More detailed information can be accessed from REPROCELL guidelines.

### Sequencing of *MTNR1B* alleles from iPSC and hESC

DNA was extracted using DNeasy Blood & Tissue Kit (Qiagen, 69506) or QuickExtract^TM^ DNA Extraction Solution (Lucigen, QE09050). A 601 bp genomic region (chr11:92,975,259- 92,975,860, genome build hg38), including the rs10830963 locus was amplified by the primers 5’-TCCAAGTAGCAGTCAGAAGC-3’ and 5’-CCAAGTGACATCTCAATGAG-3’; AmpliTaq Gold Master Mix 360 (ThermoFisher, 4398881) at recommended cycling settings (1 cycle 95 °C for 10 min, then 40 cycles of 95 °C for 15 seconds, 58 °C for 30 seconds, 72 °C for 45 seconds, then 1 cycle 72 °C for 7 minutes) was used. PCR amplicons were purified using GeneJET PCR Purification Kit (ThermoFisher, K0701) and Sanger-sequenced using either the forward 5’-GTAGCAGTCAGAAGCTGTGGTC-3’ or the reverse 5’- GCCTTCCAGAGCCTTTGTTCAG-3’ sequencing primer.

### Single-base genome editing of rs10830963 using CRISPR/Cas9 in iPSC and hESC

Cas9 protein Alt-R v3 (1072544, HDR enhancer, electroporation enhancer 1075916), as well as custom-made sgRNA and ssDNA were from Integrated DNA Technologies (Coralville, Iowa, USA). To target the risk G-allele and the non-risk C-allele, the spacer sequences TACTGGTTCTGGATAGCAGA and TACTGGTTCTGGATAGGAGA, respectively, were used in single guide RNAs. The synthetic ssDNA homology-directed repair donor template for the G-to-C allele editing was 5’-GCAGAATATTCCCATCAGGAACCTCCCAGGCAGTTACTGGTTCTGGATAGGAGATGGTGTGAATTCTTAGCATCACTGGGGGCCTGGAGGAGGGGCAGCT-3’, and for the C-to-G allele editing 5’-GCAGAATATTCCCATCAGGAACCTCCCAGGCAGTTACTGGTTCTGGATAGCAGATGGTGTGAATTCTTAGCATCACTGGGGGCCTGGAGGAGGGGCAGCT-3’. To perform the allele editing, 135 pmol Cas9 protein Alt-R v3 were incubated with 150 pmol sgRNA for 30 min before adding 200 pmol ssDNA donor template and 400 pmol electroporation enhancer. The reagents were mixed with 1 million cells resuspended in 100 μl human stem cell buffer kit 2 (Lonza, VCA-1005); the cells were electroporated using the Amaxa nucleofector IIb program B-016. The cells were then placed in culture medium supplemented with 30 μM HDR enhancer and incubated at 32 °C for two days before moving to 37 °C. After five days, an aliquot of cells was used to extract the DNA by QuickExtract^TM^ DNA Extraction Solution (Lucigen, QE09050). To assess the genomic editing efficiency, PCR amplicons were generated and sequenced as described above. Then, cells were seeded at low density, 500 cells per 10 cm dish, to generate single-cell clones. Clonal DNA was extracted and PCR of DNA sequence comprising rs10830963 was performed as above. PCR products were screened for allele editing using BsaXI restriction enzyme digests (the C allele creates a BsaXI restriction site), and the edited clones’ PCR amplicons were Sanger-sequenced to confirm proper allele editing and absence of indels.

### Culture, maintenance and freezing of iPSC and hESC

Prior to differentiation, stem cells were 2D cultured in laminin-coated (1:20; BioLamina, LN521-05) T25 flasks with Essential 8 complete medium (for iPSCs, Life Technologies A1517001 and Nutristem Media (for ESCs, Nordic Diagnostica, 05-100-1A). Thawing frozen cells required media to be supplemented with RevitaCell supplement (1:100; ThermoFisher, A2644501) for the 1^st^ 24 h of culture. Upon 80% confluency, cells were washed with DPBS-/- (ThermoFisher, 14040117) and dissociated into single cells using TrypLE select (ThermoFisher, 12563011) for 5 min at 37°C in incubator. Rho kinases inhibitor (10μM) (Sigma-Aldrich, Y-27632A, CAS 146986-50-7) was added after every round of passaging for the first 24 h of culture. For freezing of cell stocks, dissociated cells were resuspended in the appropriate volume of PSC cryopreservation medium (Gibco, A2644601) and subsequently stored at -80°C for a day and then transferred to a liquid nitrogen tank for long term storage.

### 2D differentiation of hiPSC/hESC to β-cells

Cells (⁓100,000 per well) were plated on laminin-coated (BioLamina, LN521-05) 12-well cell culture plates. Differentiation commenced according to a modified protocol [13] and was initiated when cell culture reached ⁓ 95% confluency, and the current stage was designated as differentiation day 0. For day 0-5, all cells were cultured in RPMI (ThermoFisher, 61870-044) medium; day 5-50 cells were cultured in DMEM-F12 (ThermoFisher, 31331-093). The cell media (1ml/well) were supplemented with factors at each developmental stage as follows. Day 0-1, Activin A (100ng/ml) (Peprotech, 120-14E), CHIR99021 (3μM) (Sigma Aldrich, SML1046-5MG). Day 1-5, Activin A (100ng/ml) and B27- insulin (ThermoFisher, A1895601). Day 5-8, Retinoic Acid (2μM) (Sigma Aldrich, R2625-50MG). Day 8-11, FGF2 (64 ng/ml) (Peprotech, 100-18B). Day 11-14, TBP (0.5μM) (ThermoFisher, 4351370) and Noggin (100 ng/ml) (Peprotech, 120-10C). Day 15-17, Forskolin (10μM) (Sigma Aldrich, f3917-25mg), Alk5i (4.5μM) (SantaCruz, sc-221234A), Noggin (100 ng/ml) (Peprotech, 120-10C), Nicotinamide (10mM) (Sigma Aldrich, N0636-100G). Day 17-30 Forskolin (10μM) (Sigma Aldrich, f3917-25mg), Alk5i (4.5μM) (SantaCruz, sc-221234A), Noggin (100 ng/ml) (Peprotech, 120-10C), Nicotinamide (10mM) (Sigma Aldrich, N0636-100G). In addition, Day 5-50 differentiation media were supplemented with B27 (ThermoFisher, 17504044). Cell culture medium was replaced every 24 hours from days 0-17 and every 48 hours from days 17- 50.

### Immunoblotting

Cell extracts were prepared using RIPA buffer (Sigma, R0278) supplemented with Pierce Protease Inhibitor Cocktail (ThermoFisher, A32963); the protein concentrations were quantified by BCA protein assay (ThermoFisher, 23225). 20 μg protein were run on a 4-20% reducing SDS-PAGE Mini-PROTEAN TGX gel (BioRad, 4568094), and transferred to a PVDF membrane (BioRad, 1704157) that was blocked for 1 h in 5% BSA (Sigma, A4503) in TBS (Biorad, 170-6435). Next, the membrane was immunoblotted for 16 hours at 4 °C in blotting buffer (3% BSA in TBS with 0.1 % Tween-20 (Sigma, P7949) with 1 μg/ml rabbit anti-MTNR1B (ThermoFisher, PA5-102107), washed 4 x 5 minutes in washing buffer (TBS with 0.1% Tween-20), then incubated for 1 h at room temperature with 0.1 μg/ml HRP-conjugated goat anti-rabbit (BioRad, 162-0177) in blotting buffer, washed 4 x 5 minutes with washing buffer, then developed using Clarity Western ECL substrate (BioRad, 1705060) and a CCD camera. For reprobing, the membrane was stripped with Restore Western Blotting Stripping Buffer (ThermoFisher, PIER21059), and the procedure was repeated as above but using 0.2 μg/ml mouse anti-α-tubulin (Abcam 7291) as primary, and 0.1 μg/ml HRP-conjugated goat anti-mouse antibody (BioRad, 1706516) as secondary antibody. The bands on the immunoblots were quantified using ImageJ software (NIH). MTNR1B signal was normalized to β-tubulin signal.

### Immunocytochemistry

hiPSC, hESCs and fully differentiated cells were fixed directly on the culture plates with 4% paraformaldehyde (PFA, Histolab, 2176) for 30 min at room temperature (RT), permeabilized with 0.1% triton X-100 (Sigma) in PBS (Gibco) for 15 min at RT and blocked with blocking buffer (5% normal donkey serum (NDS) (Abcam, ab7475) + 1% bovine serum albumin (BSA, Sigma, 05482) + 0.1% Tween-20 (Sigma, P7949) in PBS (Gibco, 18912014) overnight at RT. Primary and secondary antibodies used in this study are described in Supplementary Table 1. Primary antibody incubation was overnight at 4°C followed by incubation with secondary antibodies in blocking buffer overnight at RT (1:1000, ThermoFisher). Nuclei were stained with 4′,6-diamidino-2-phenylindole, dihydrochloride (DAPI, 1:2000, ThermoFisher, 62248) in 1% BSA in PBS for 10 min at RT. Cells were covered with fluorescence mounting medium (DAKO, S3023) and stored at 4°C. Images were captured by a laser scanning confocal microscope (LSM780, Zeiss). Final images were processed and compiled using Adobe Photoshop and InDesign. Antibodies are described in Supplementary table 1.

### qRT-PCR on iPSC and hESC during differentiation stages

RNA at the defined stages of differentiation (day 0, 1, 5, 8, 11, 14, 17, 27, 35, 50) was extracted using RNeasy Plus mini kit (Qiagen, 74136) according to the manufacturer’s guidelines. cDNA synthesis with 500 ng total RNA (bulk cells) was performed using RevertAid first strand cDNA synthesis kit (ThermoFisher, K1622) according to the manufacturer’s guidelines. Real-time qPCR reactions were set up in duplicates for each sample, using Taqman assays and Taqman Universal master mix (ThermoFisher, 4305719) in real-time qPCR system (Applied Biosystems, Quant Studio 7 Flex) using StepOnePlus software. qRT-PCR data were normalized to the geometric mean of two housekeeping genes (*TBP*, TATA-box binding protein) and *PPIA* (Cyclophilin A), using the delta C_T_ method. All TaqMan assays used in this study are listed in Supplementary Table 2.

### pLenti-HIP-GFP virus production

Lentiviral vectors were generated, as previously described [14], with titers ranging from 10^7^ to 10^8^ TU/ml and determined by fluorescence-activated sorting (FACS). Briefly, HEK293T cells were cultured to reach a confluency of 80-90% on the day of transfection. For the production, third-generation packaging, and envelope vectors (pMDL, psRev, and pMD2G) were used in conjunction with Polyethyleneimine (PEI Polysciences PN 23966) in DPBS (Sigma Aldrich, D8537). The supernatant was then collected, filtered, and centrifuged at 25,000g for 1.5 hours at 4°C. The supernatant was removed from the tubes and the virus was resuspended in PBS (Sigma Aldrich, P4474), and left at 4°C. The resulting lentivirus was aliquoted and stored at −80°C. Generated lentivirus was also tested and titrated in human EndoC-βH1 line (Supplementary figure 4).

### Fluorescence-activated cell sorting

At day 45 of differentiation, cells were infected with pLenti-HIP-GFP virus (MOI=2). Five days post-transduction, robust GFP green fluorescence appeared in β-cells. For FACS, single cells were prepared using TrypLE select (ThermoFisher, 12563011). All washes were performed in FACS buffer (2% FBS (Sigma Aldrich, A7030-100G) in DPBS-/- (ThermoFisher, 14040117). Cellular debris were removed as described by the manufacturer’s guidelines (Miltenyibiotec, 130-109-398). Cells were filtered through 30uM filter (BD Biosciences, 340627) and sorted (BD Aria Fusion) for GFP^+^ cells (Supplementary figure 5) in Eppendorf tubes containing FACS buffer. Cells were redistributed in lysis buffer (Norgen Biotech 51800) and stored in -80°C.

### qRT-PCR on sorted β-cells

Sorted, GFP**-**expressing β-cells were subsequently processed to obtain RNA, using Single Cell RNA Purification Kit (Norgen Biotech 51800). RNA was quantified by nanodrop spectrophotometer (ND-1000). cDNA was synthesized from 50-80 ng of total RNA according to manufacturer’s protocols using SuperScipt IV VILO Mastermix with ezDNASE Enzyme kit (ThermoFisher, <u>11756050</u>). Real-time qPCR reactions were set up in duplicates for each sample using Taqman assays and Taqman Fast Advanced master mix (ThermoFisher, 4444556) in real-time qPCR system (Applied Biosystems, Quant Studio 7 Flex) using StepOnePlus software. qRT-PCR data were normalized to the geometric mean of two housekeeping genes (*TBP* and *PPIA* using the delta C_T_ method. All TaqMan assays used in this study are listed in Supplementary Table 2.

### Insulin secretion

Differentiated β-cells cultured until day 50-60 in 12-well plates were starved in secretion assay buffer (SAB; 114 mmol/l NaCl, 4.7 mmol/l KCl, 1.2 mmol/l KH_2_PO_4_, 1.16 mmol/l MgSO_4_, 25.5 mmol/l NaHCO_3_, 20 mmol/l HEPES, 2.5 mmol/l CaCl_2_ and 0.2% BSA (fatty acid free), pH 7.2) supplemented with 1 mM glucose at 37 °C for 2 hours. Following starvation, cells were stimulated for 1 hour with SAB containing either low glucose (LG, 1 mM), high glucose (HG, 20 mM) or HG + IBMX (50 μM or 100 μM). Each of the four treatments was performed in the presence or absence of 100 nM melatonin (Sigma, M5250). Supernatant was collected from the wells and centrifuged (4000 x g, 5 min, 4 °C). To quantitate intracellular insulin and protein, cells were rinsed in the wells with DBPS and lysed with ice-cold RIPA buffer supplemented with protease/phosphatase inhibitor (ThermoFisher, A32961), scraped from the well bottom and rotated on a shaker (1600 rpm, 30 min, 6 °C). Lysates were centrifuged (12000 x g, 15 min, 4 °C) and supernatant collected. Insulin was quantified using human insulin ELISA kit (Mercodia, 10-1113-10) according to the manufacturer’s guidelines. Insulin concentration in the samples was interpolated from the standard curve using cubic spline regression (Prism/GraphPad). Protein concentration was determined with Pierce™ BCA Protein Assay Kit (ThermoFisher, 23225).

### Data Analysis and interpretation

Statistical testing for two groups were performed using two-tailed Student’s *t* test and the Mann–Whitney *U* test was applied when normal distribution was not apparent. Insulin secretion measurements comparing more than three groups (C/C and G/G genotypes combined with Melatonin, IBMX and the combination thereof) was analysed wth RM One-way ANOVA with Šídák test to correct for multiple comparison, after normalization to respective glucose control of each group. A *p*-value of <0.05 was considered statistically significant. Data are presented as mean±SD. All analyses were undertaken using the Graphpad Prism 10 software.

## Results

### CRISPR-Cas9 editing of hiPSC and hESC at the *MTNR1B* locus

hiPSC from two human donors (MF002B2 and MF007C1) were subjected to single-base genome editing of the *MTNR1B* (rs10830963) risk allele (G/G) using CRISPR/Cas9 (See Table 1a-c for patient information and Fig. 1a for work-flow of skin biopsy procurements to create hiPSC). Hereby, isogenic cell lines were generated carrying non-risk variants (C/C); editing efficiency was >90% (Fig. 2a-c). In addition, the hESC line, HUES4 [15], was edited at the *MTNR1B* locus (original genotype C/G) to either C/C (non-risk) or G/G (risk), respectively (Fig. 2d), with an efficiency comparable to that in hiPSC clones. Successful editing was confirmed by BsaXI enzyme restriction digestion, where cutting of DNA with G/G risk genotype generated a 510 bp DNA band while DNA with C/C non-risk genotype generated a 210 bp band (Fig. 2e). Prior to differentiation of hiPSC and hESC to β-cells, MTNR1B protein was readily detected in all cultures (Supplementary Fig. 2).

**Figure 1.**
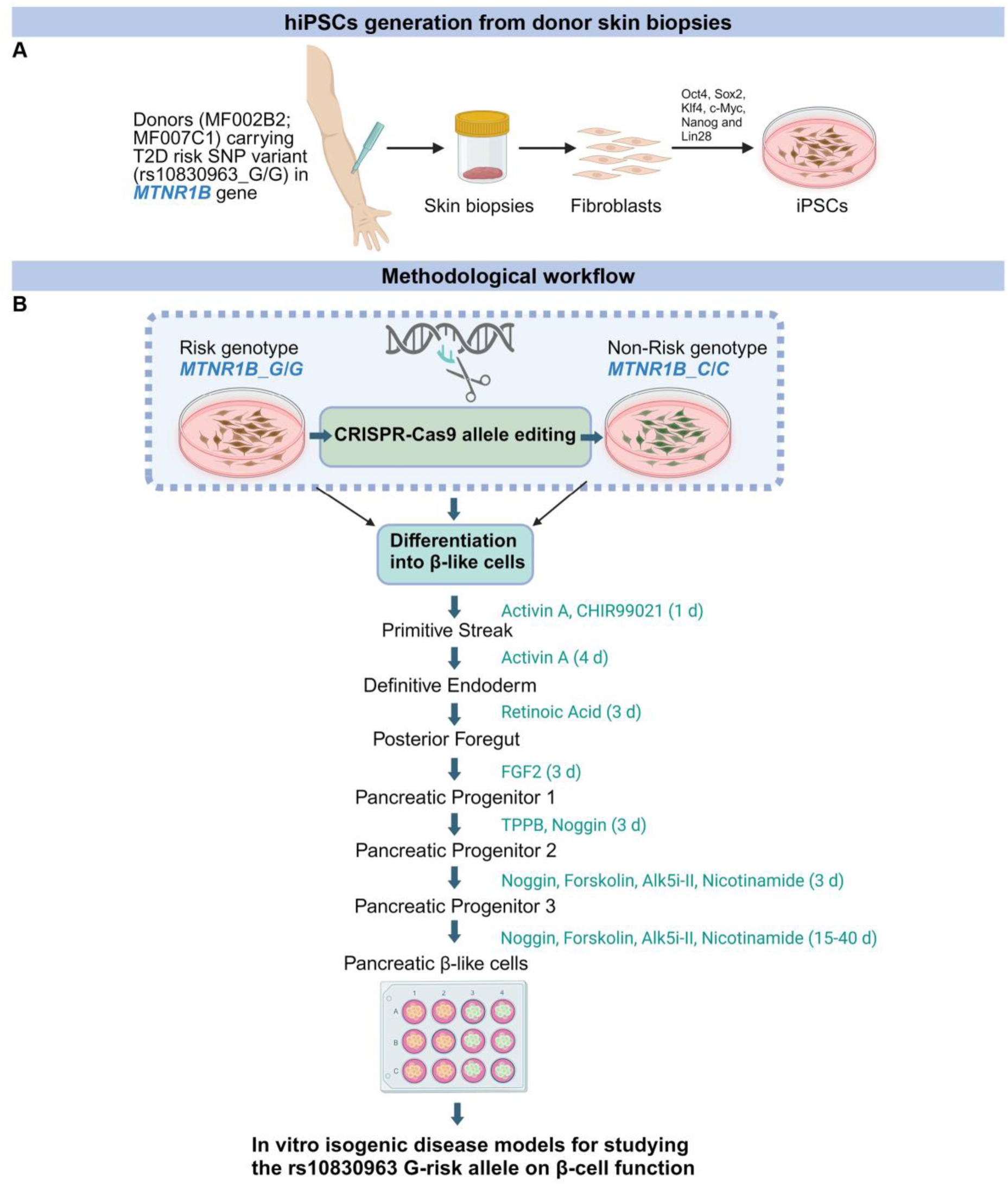
Graphical representation of derivation of iPSCs and reprogramming into β- cells *in vitro*. (A, B) The image illustrates the workflow of procuring skin biopsies from patients, iPSC reprogramming, single base genome editing, and differentiation into β-cells. (B) The procedure creates two isogenic cell lines, which differ only with respect to one single base, i.e., the single nucleotide polymorphism (SNP) that constitutes the risk allele.

**Figure 2.**
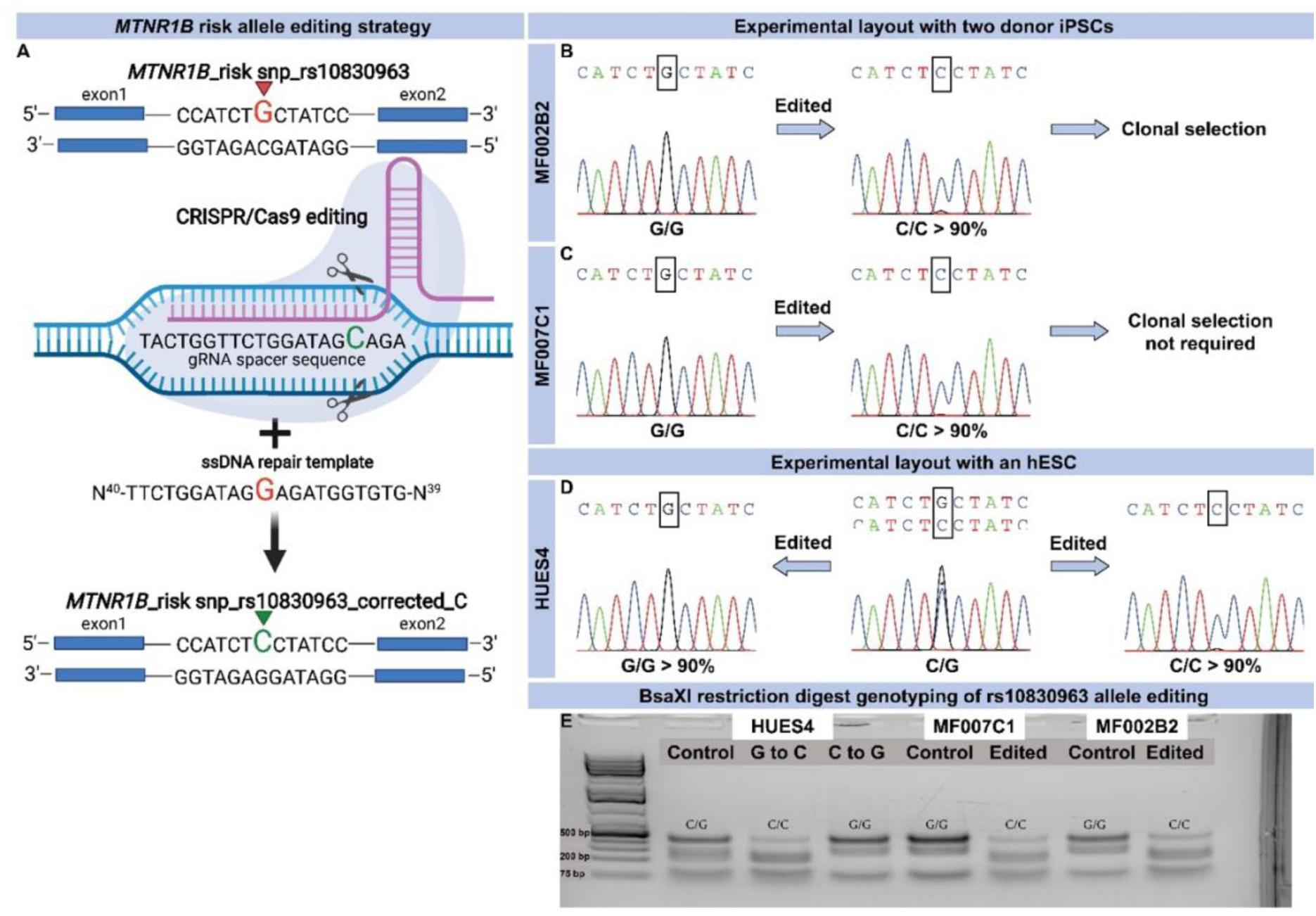
Genome editing of the *MTNR1B* T2D risk allele and edited clone selection strategy. (A) The graphical representation shows the sequence surrounding rs10830963 in the intron of the *MTNR1B gene*. G, indicated by red arrowhead, is the minor allele conferring the increased risk of impaired insulin secretion and future risk of T2D. A guide RNA (gRNA; purple) targeting Cas9 to this sequence was prepared and introduced into hiPSC along with a single stranded (ss) DNA template. It contains a C instead of G (red; NB – G will be transcribed into C); after Cas9 breaks the chain, repair will introduce C (green) instead of G, resulting in a “correction” of the risk G-allele in edited cells. (B-C) Sanger sequencing of DNA in iPSCs carrying the homozygous (G/G) risk allele of *MTNR1B* and upon editing, sequencing after single base gene editing in a pool of hiPSC where the vast majority of cells now carries the non-risk C-allele. A small peak below the major peak indicates that a minor fraction of cells still remains unedited. (B) MF002B2 edited cells were expanded and clonal cell lines were selected as pure edited and control lines. (C) As the editing efficiency was more than 90%, clonal selection for edited MF007C1 cells was not performed. (D) hESCs (HUES4) are of the heterozygous G/C genotype at the rs10830963 locus in *MTNR1B*. Therefore, these cells were edited both ways from G to C and C to G in parallel to create cell lines carrying homozygous non-risk C/C and risk G/G risk alleles of *MTNR1B.* Editing efficiency was more than 90%, hence no clonal cell lines were prepared. (E) Editing was also assessed by BsaXI restriction digestion where G/G risk genotype resulted in a prominent DNA band at a roughly 510 bp size and C/C non-risk genotype at a roughly 210 bp size.

**Table 1:** Donor information, *MTNR1B* genotype and their diabetogenic profile. (A) Table exhibits the two donors included in the study and their risk SNP G/G genotype at the *MTNR1B* locus. (B) Shows insulin and (C) glucose measurements upon oral glucose tolerance tests for both donors carrying the risk allele.

| Donor ID | AGE | SEX | BMI | Fasting Glucose | MTNR1B genotype rs10830963 |
| --- | --- | --- | --- | --- | --- |
| MF002 | 71 | Male | 27,53 | 7,60 | G/G risk |
| MF007 | 63 | Male | 36,09 | 9,50 | G/G risk |

B.
| Donor ID | Insulin, 0 min | Insulin, 15 min | Insulin, 30 min | Insulin, 45 min | Insulin, 60 min | Insulin, 90 min | Insulin, 120 min |
| --- | --- | --- | --- | --- | --- | --- | --- |
| MF002 | 15,0 | 24,0 | 32,0 | 29,0 | 31,0 | 50,0 | 63,0 |
| MF007 | 26,0 | 43,0 | 63,0 | 69,0 | 71,0 | 88,0 | 104,0 |

C.
| Donor ID | Glucose, 0 min | Glucose, 15 min | Glucose, 30 min | Glucose, 45 min | Glucose, 60 min | Glucose, 90 min | Glucose, 120 min |
| --- | --- | --- | --- | --- | --- | --- | --- |
| MF002 | 7,5 | 8,2 | 9,4 | 8,8 | 11,5 | 14,6 | 15,5 |
| MF007 | 9,4 | 10,1 | 13,3 | 15,7 | 18,5 | 19,4 | 19,5 |

### Differentiation of hiPSC and hESC into β-cells

After successful editing, hiPSC and hESC clones were differentiated into β-cells employing a protocol for 2D-culture [13], where addition of small molecules and growth factors enabled pancreatic and islet cell development (Fig. 1b). All clones were transcriptionally and immunohistochemically evaluated following differentiation (day 0-50) with regards to pancreatic endocrine markers (PDX1 and NKX6.1) as well as for insulin (C-peptide) during the later stages of differentiation (Fig. 3a-d for hiPSCs; e-h for hESCs). Importantly, insulin was expressed in both hiPSC and hESCs from day 35 (Fig. 3i and k). This could also be confirmed at the protein level by immunohistochemical staining of hiPSC and hESCs with antibodies to C-peptide at day 50 of differentiation (Fig. 3a and e). Of note, insulin gene (*INS*) expression was approximately 20-fold higher in the hESCs as compared to hiPSCs (Fig. 3k vs. 3i), suggesting an enhanced capacity of hESCs to differentiate into β-cells. Similarly, *PDX1* expression started at day 6 of differentiation in both hiPSC and hESC: its expression varied during the course of differentiation; hESCs expressed *PDX1* at a 10-fold higher level at the end of the protocol (Fig. 3l vs. 3j). Insulin expression (C-peptide staining) in cell cultures was evident throughout the culture (Supplementary Fig. 3 - tile scans; a and b, hiPSC clones and c, hESCs). Immunohistochemical staining revealed both insulin- and glucagon-positive cells, with some bi-hormonal cells observed in the cell cultures (Supplementary Fig. 3d-f). Glucagon (*GCG*) gene expression in hiPSCs and hESCs was detected from day 35 in hiPSC and hESCs (Supplementary fig. 3g and h respectively). Importantly, *GCG* gene expression was approximately 18-fold lower than *INS* gene expression in hiPSC derived endocrine cell cultures and 10-fold lower in in hESC derived endocrine cell cultures.

**Figure 3.**
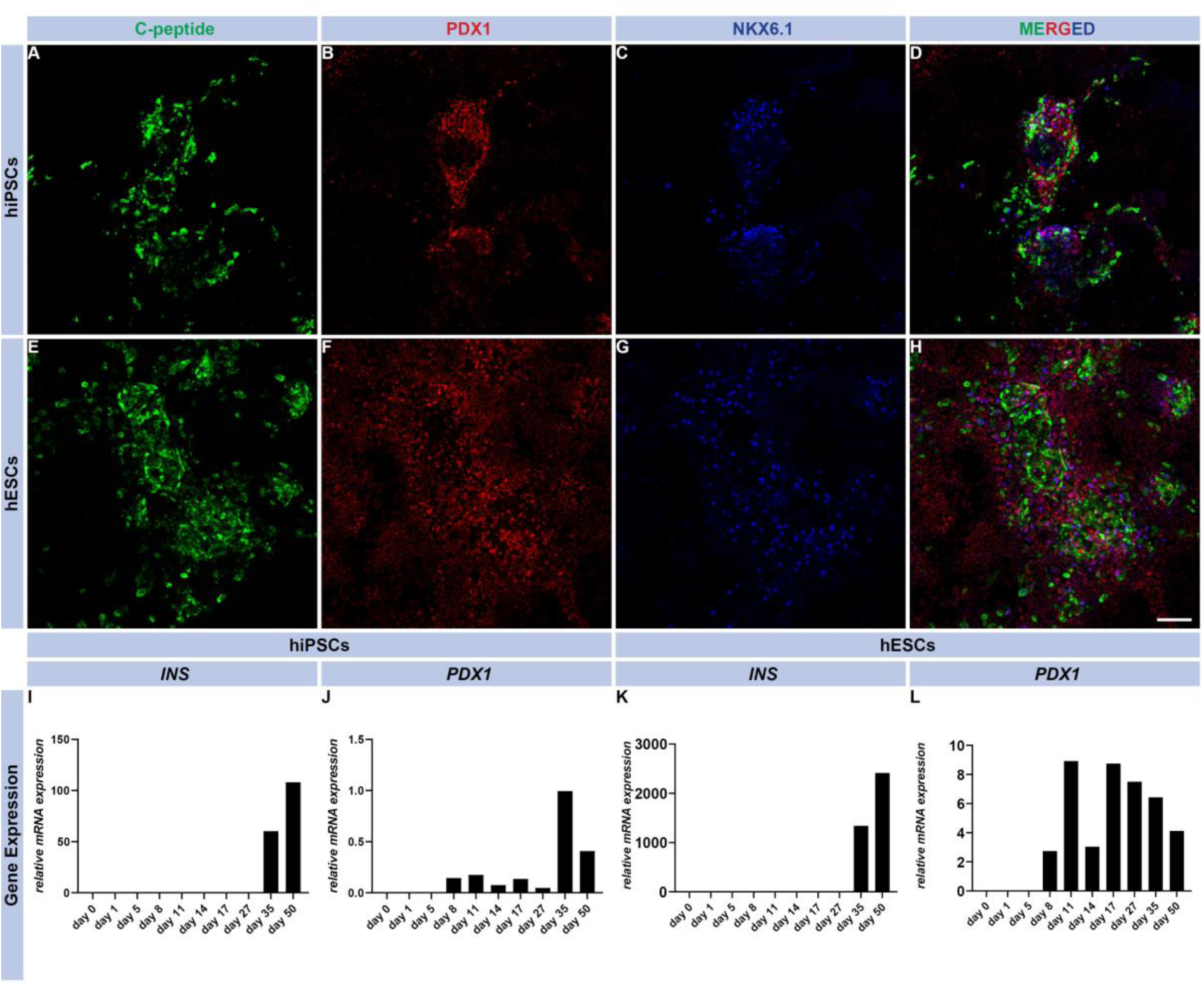
hiPSC and hESC lines and β-cell differentiation characterization. (A-H) Immunocytochemistry images of differentiated hiPSCs and hESCs that display C-peptide^+^ cells (β-cells in green) co-expressing PDX1 (red), and NKX6.1 (blue) in fully differentiated cells at day 50 of differentiation. The scale bar represents 100 μm, and the images were captured at 20X magnification with a confocal microscope. (I-L) Gene expression profile of *INS* and *PDX1* in β-cells differentiated for 50 days.

### The *MTNR1B* risk variant rs10830963 is associated with altered protein but not mRNA levels in hiPSC differentiated to β-cells

To determine whether the *MTNR1B* risk and non-risk variant impacted expression of *MTNR1B*, we used isogenic clones of hiPSC carrying either edited (C/C; non-risk) and non-edited (G/G; risk) *MTNR1B* genotypes (Fig 4A). Six clones of each genotype (C/C and G/G) were chosen, using 2D-cultures for subsequent differentiation of hiPSC into β-cells. At day 50 of differentiation, expression of *INS* and *MTNR1B* was determined by qPCR (Fig. 4b and c). No difference in *INS* or *MTNR1B* expression was observed between the genotypes (6 clones/genotype from one patient donor). At the protein level, however, western blotting demonstrated a trending increase (p=0.09) of MTNR1B protein in β-cells carrying the G/G risk allele (Fig, 4e).

**Figure 4.**
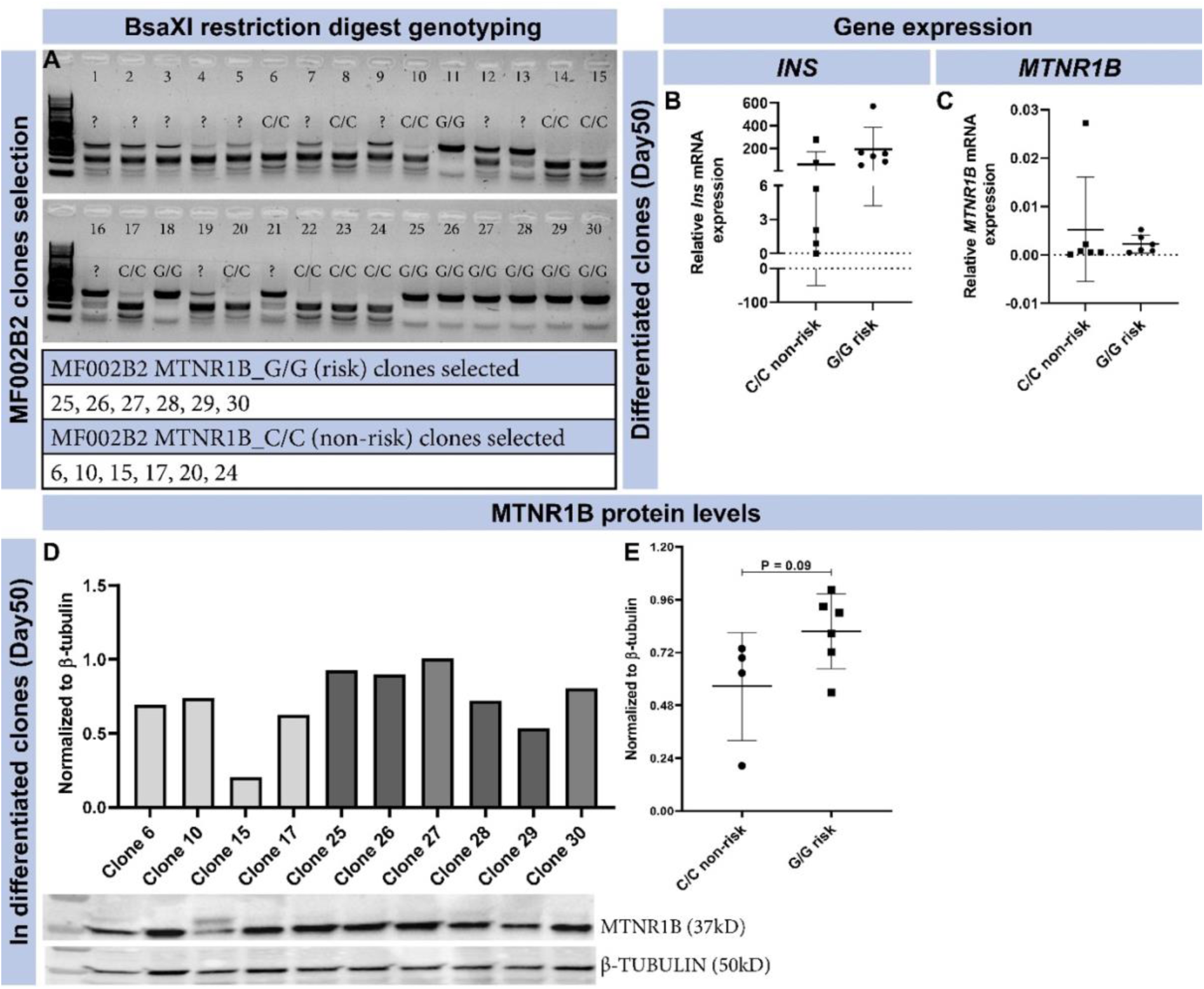
MF002B2 hiPSC line, clonal selection and *MTNR1B* expression. (A) Edited non-risk (C/C) and risk (G/G) clones were selected based on BsaXI restriction digestion genotyping. All clones with clear bands were selected (lower panel in A). (B, C) At the end of differentiation, cells (bulk) were assessed for gene expression of *INS* (B) and *MTNR1B* (C) (N=6, mean ± SD; each data point represents one individual differentiation experiment from each selected clone). (D-E) MTNR1B protein levels were assessed at the end of differentiation in bulk cells and presented as individual bars for each clone (D) as well as cumulatively (E) (N=4-6, mean ± SD; each data point represents one individual differentiation experiment from each selected clone, (p>0.05 was considered significant).

To improve our ability to detect differences in *MTNR1B* expression in β-cells from isogenic risk and non-risk allele carriers, we further increased the proportion of β-cells by FACS of edited and non-edited hiPSC and hESCs. Five days prior to FACS, cell lines were infected with a pLenti vector expressing green florescent protein (GFP) under the control of the human insulin promoter (HIP; MOI 2; Fig 5a-c). Hereby, insulin-producing cells would be detected and amenable to sorting by FACS (Suppl. Fig. 5). Next, we investigated the expression of *INS* and *MTNR1B* in these cell preparations (Fig 5a-g). Again, we were unable to detect any differences in mRNA expression of either *INS* or *MTNR1B* in the β-cells derived from hiPSC with the G/G and C/C genotypes, respectively. Interestingly, we observed reduced *MTNR1B* gene expression (P=0.02) in β-cells derived from hESCs carrying the G/G risk allele compared with hESCs of the C/C genotype (Fig 5 i). This is contrary to what we previously have observed in human islets of Langerhans, utilizing qPCR or RNA sequencing respectively [2, 16], where a significant increase in the *MTNR1B* mRNA in non-diabetic G/G risk allele carriers was evident. Regardless of this and of which cell line analyzed, *MTNR1B* expression was consistently very low, just above the level of detection.

**Figure 5.**
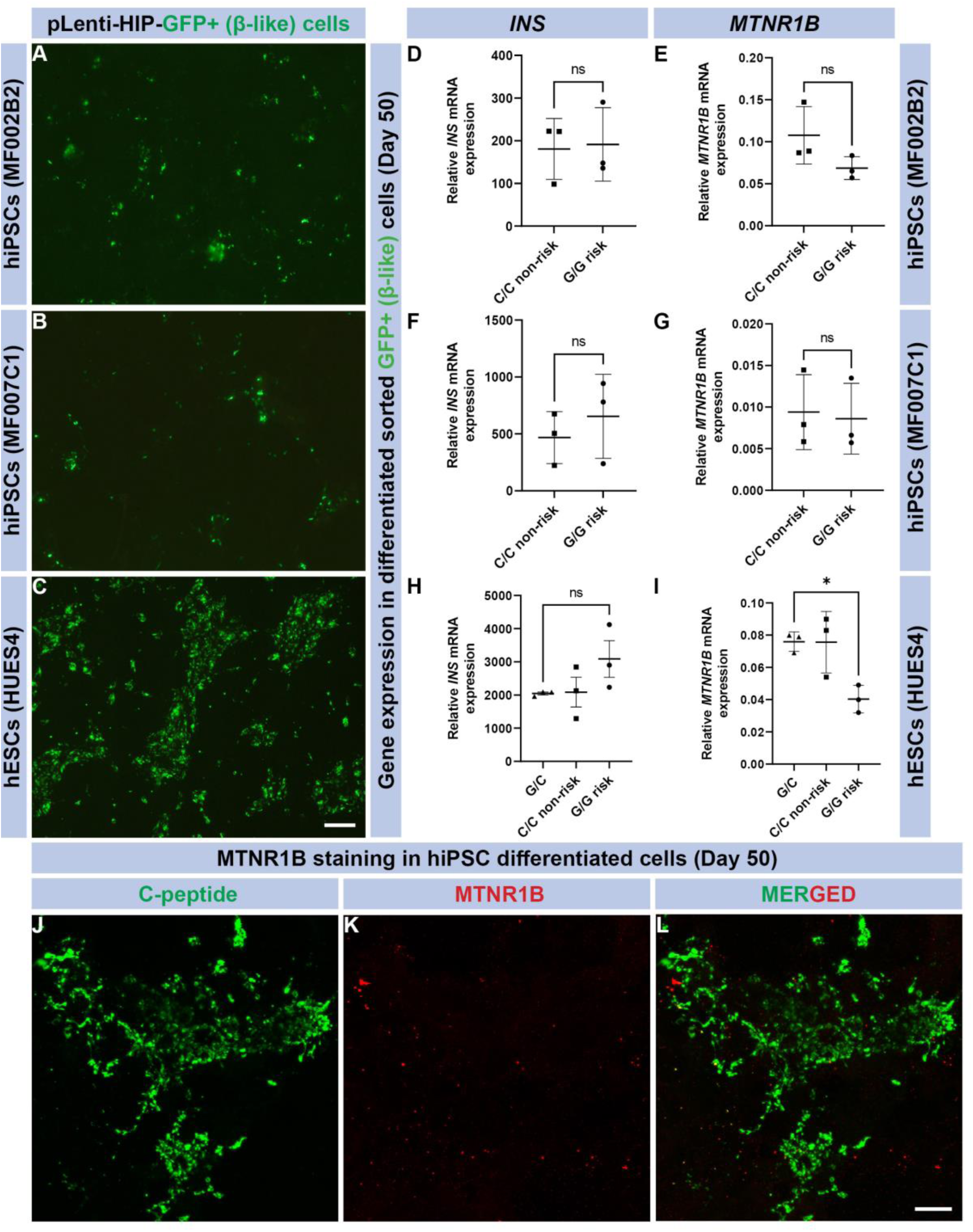
*MTNR1B* expression in fully differentiated β-cells. (A-C) Images of GFP expression (green) in β-cells post lentivirus infection. (D-I) Graphs show *INS* (D, F, H) and *MTNR1B* (E,G,I) expression in sorted GFP+ β-cells (N=3, mean ± SD; each data point represents one individual differentiation experiment). Unpaired t-test with Welch’s correction was performed. (J-L) Immunocytochemistry images show robust C-Peptide+ cells (β-cells, green) with absence of MTNR1B (red) co-expression. The scale bar represents 100 μm, and the images were captured at 10X magnification with a confocal microscope.

### Insulin secretion from β-cells derived from hiPSC and hESC

To determine functional changes in isogenic cell lines upon editing of the *MTNR1B* G-risk allele, we exposed β-cells (derived from both hiPSC and hESC) to either low (LG 1 mM) or high (HG 20 mM) glucose, and 3-isobutyl-1-methylxanthine (IBMX 50 or 100 μM) with or without melatonin (100 nM; Figure 6 a-h). Neither hiPSC- nor hESC-derived β-cells responded with an increase in insulin secretion to a rise in glucose concentration (Fig. 6 a and e; shown as a fold to control (LG)) for C/C and G/G genotypes. Insulin secretion appeared to increase in both 50 and 100 μM of IBMX (Fig. 6c) in hiPSCs and hESCs (Fig. 6g), however, this did not reach statistical significance. In addition, addition of melatonin significantly reduced insulin secretion in hiPSC-derived β-cells carrying the G/G genotype (as compared to C/C) at LG concentrations (*p*=0.015; Fig. 6b). Moreover, the combination of 50 μM IBMX and melatonin reduced insulin release from hESC-derived β-cells carrying the G-allele as compared to C-allele (Fig. 6d; *p*=0.08).

**Figure 6.**
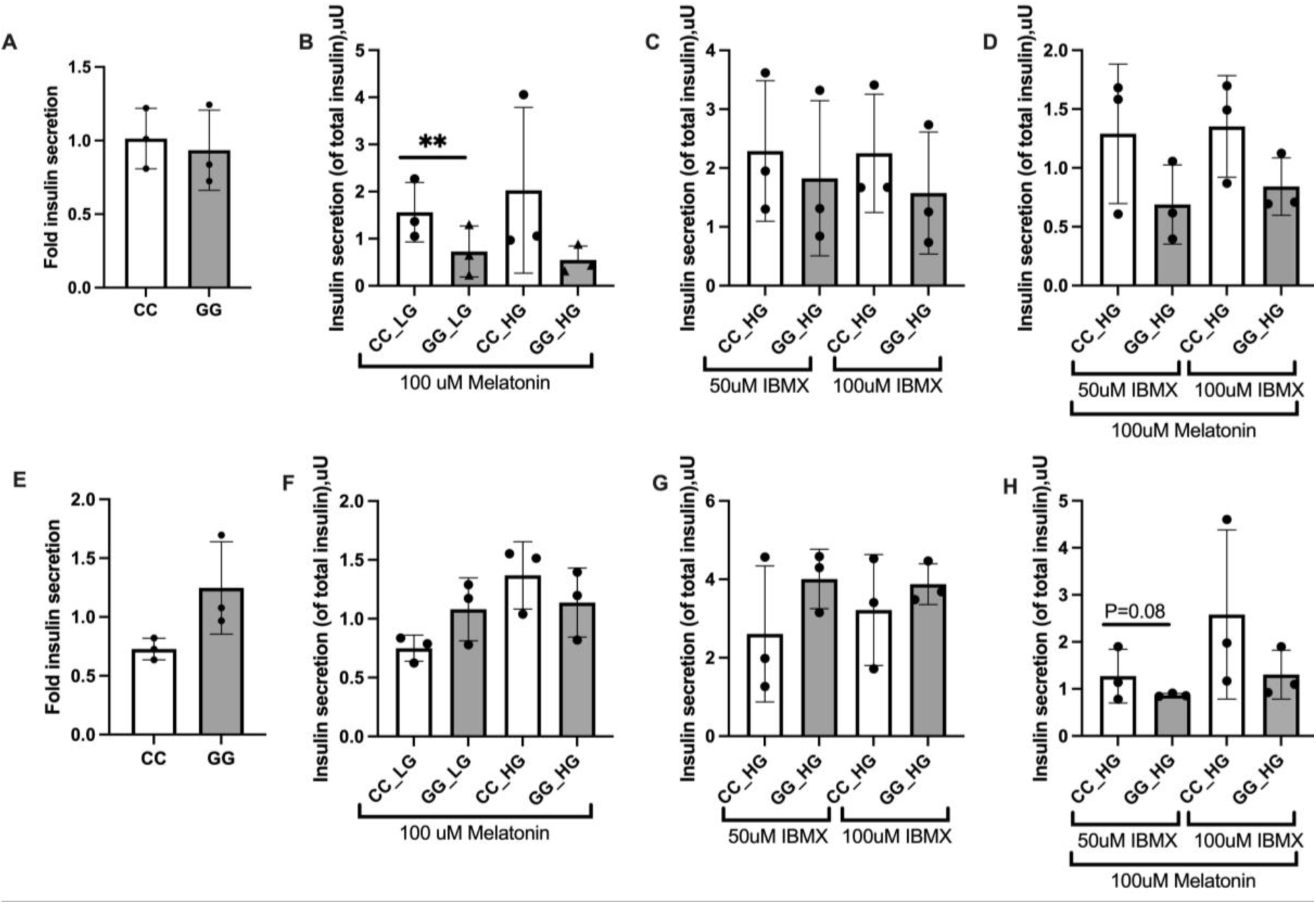
Insulin secretion in differentiated cells. Secreted insulin in hiPSC derived β-cells (A-D), measured after stimulation for 1 h with 1 mM glucose (LG), 20 mM glucose (HG) (A), stimulation with LG, HG and Melatonin (100nM) (B). Stimulation with HG and 50 µM or 100 µM, IBMX (C). Stimulation with HG and 50 µM or 100 µM IBMX with Melatonin (100nM) (D). Secreted insulin in hESC derived β-cells (E-H), measured after stimulation for 1 h with LG, and 20 HG (E), stimulation with LG, HG and Melatonin (100nM) (F). Stimulation with HG and 50 µM or 100 µM, IBMX (G). Stimulation with HG and 50 µM or 100 µM, IBMX with Melatonin (100nM) (H). (N=3 biological replicates, mean ± SD) *p*<0.05*.

## Discussion

Molecular mechanisms linked to genetic signals identified by GWAS could help elucidate the causes of β-cell dysfunction in T2D. However, functional consequences of these signals in β- cells presently remain unknown. Here, we describe a stem cell-based approach, utilizing genome editing of hiPSC and hESCs followed by differentiation into b-cells, to elucidate functional effects risk alleles. Specifically, we targeted the SNP rs10830963 mapping to an intron of the gene encoding the MTNR1B receptor, which is a robust risk allele for T2D [2, 6–9].

We performed differentiation of hiPSC and hESC, utilizing a previously established 2D-based protocol [13], permitting differentiation of stem cells to β-cells. Both hiPSCs and hESCs differentiated accordingly, with the expected rising expression of endocrine markers, followed by increasing expression of *INS* and *GCG* at mRNA and protein levels during the later stages of differentiation, with *INS* expression being several fold higher at the termination of the protocol in both hiPSCs and hESCs.

Prior to differentiation of hiPSCs and hESCs, we utilized CRISPR/Cas9 to perform single-base genome editing. In hiPSCs, we edited the risk allele (G) to non-risk (C) to generate isogenic cell lines, with genotypes G/G and C/C. Utilizing isogenic cell lines for comparison is essential as genetic heterogeneity between humans is considerable. Ideally, isogenic cell lines will differ only with regard to the edited SNP (from G to C). We also edited the DNA of hESCs cells, which initially were heterozygous (C/G) at the rs10830963 locus: we edited this allele to either the non-risk (C/C) or risk (G/G) genotype. High efficiency (> 90%) and fidelity (minimal indels) were achieved for all cell clones. Previous studies have shown limitations of the CRISPR/Cas9 system in its applicability due to, e.g., low transfection efficiency, toxicity, difficulty in obtaining pure clones of edited cells, and off-target effects [17]. However, here, no issues with editing were observed; single-base edited hiPSCs and hESCs were successfully differentiated into b-cells.

We have previously shown that the *MTNR1B* rs10830963 SNP is an eQTL in human islets of Langerhans [2]. In addition, there is evidence that this SNP may influence gene expression [11], as the SNP has been shown to increase FOXA2-bound enhancer activity in islet- and liver-derived cells. In addition to this, allele-specific differences of the transcription factor NEUROD1 binding in islet-derived cells have been observed [11]. This suggests that the rs10830963 SNP confers increased islet or b-cell *MTNR1B* expression, thus constituting an eQTL. Demonstrating cellular or molecular proof of function of rs10830963 is critical, as it could merely represent a tag SNP in the haploblock, where additional SNPs are present, and may influence gene expression and cellular function, either separately or combined. Presently, causality of rs10830963 can only be determined by specifically editing this SNP in a relevant model and determine functional consequences (e.g., prove that the SNP is an eQTL that increases the *MTNR1B* expression in carriers of the G allele). Indeed, this may prove to be difficult, as the effect size of a single SNP may be relatively small, given that multiple genetic signals associate with glycaemic traits and insulin secretion in T2D.

mRNA expression of *MTNR1B* in risk (G/G) and non-risk isogenic clones (both hiPSCs and hESCs) was determined. In general, the expression of *MTNR1B* was very low, both in sorted and unsorted β-cells. The inability to detect and determine the expression of *MTNR1B* mRNA and protein in islet cells has previously been discussed: studies utilizing global knock out mice of *Mt1* and *Mt2* reveal immunostaining of Mt2 protein localized to β-cells, while Mt1 was observed in α-cells of the islet [18]. *MTNR1A* and *MTNR1B* protein expression and density in human pancreatic β-cells have been confirmed [2, 19, 20], but with lower *MTNR1A* and *MTNR1B* expression in α-cells [20]. Others show *MTNR1A* expression predominantly in α-cells of human islets, while low levels of *MTNR1B* expression is detected [21]. The inherently low mRNA expression of *MTNR1B* in human islets has made it difficult to determine islet cell-specific localization, even with recent efforts using single cell RNA sequencing [22]. In this context, studies on islet cell heterogeneity, genetic variation, metabolic state, half-life and stability of the *MTNR1B* protein/mRNA may provide further insight. Interestingly, *MTNR1B* expression was readily detected in isolated human islets from both risk and non-risk allele carriers, with a significant increase in expression in risk allele carriers [2] by qPCR or RNA sequencing [16]. Of note here, we readily detected MTNR1B protein, both in undifferentiated hiPSCs and hESCs with Western blot and in differentiated β-cells with Western blot and immunocytochemistry. In fact, protein expression of MTNR1B was nominally increased in β- cells from risk allele carriers (albeit not reaching full statistical significance; *p*=0.09), agreeing with the notion that the *MTNR1B* risk SNP (rs10830963) is indeed an eQTL; this assumes that mRNA levels correspond to protein levels. Again, the effect size of a single SNP alteration, may be obscured by numerous other genetic signals. Indeed, similar studies of monogenic forms of diabetes (e.g., MODY) report functional changes, however, in these cases the effects size is profoundly greater [23–25].

Consistent with other studies [26–28], our stem cell derived β-cells do not release insulin in response to an elevation of exogenously added glucose. However, we observed a significant reduction of insulin release upon addition of melatonin in hiPSCs derived β-cells carrying the G-risk allele as compared to β-cells with the C-allele. Similarily, melatonin nominally reduced IBMX-stimulated insulin secretion in hESCs derived β-cells with the G/G as compared to the C/C genotype (although borderline significant p=0.08). This suggest that hESC and hiPSC derived β-cells carrying the G-allele may be more sensitive to melatonin treatment, suggesting that the rs10830963 SNP mapping to the *MTNR1B* locus indeed represses insulin release. Importantly, melatonin has previously been shown to decrease insulin seceretion, where acute effects of melatonin on insulin release are mediated by reduced formation of cAMP with a net inhibitory effect on insulin release [29]. Stimulation of β-cells by glucose is known to increase intracellular cAMP, whereas melatonin blocks cAMP formation using various signalling pathways in clonal β-cells, rodent and human islets [30–33]. Of note, stimulatory effects of melatonin on insulin secretion under certain conditions have also been reported, particularly following prolonged exposure in rodent islets [33] and human islets [21]. Besides the well-known receptor mediated effects, melatonin can act as an antioxidant and reportedly reduces oxidative stress in β-cells *in vitro* [34, 35].

It is established that stem cell derived β-cells mature and function *in vivo,* following transplantation into mice [36], but the consistency of response *in vitro* is less robust. The reduced insulin secretory response of β-cells derived from stem cells likely emanates from lack of humoral factors, and/or formation of an effective niche, as well as of microvasculature, all of which are present *in vivo* during organogenesis, thereby promoting β-cell maturation and function. Recently, however, several new protocols have been developed with improved yield and functionality of hiPSC-derived β-cells, utilizing 3D-based approaches [37–40]. Still, a certain degree of immaturity appears to be present in all stem-cell derived β-cells [36]. One reason for this was recently explored by metabolic profiling of stem cell derived β-cells to determine their capacity to sense glucose [41]. Here, reduced anaplerotic cycling in the mitochondria was identified as a potential cause of reduced glucose-stimulated insulin secretion. Importantly, insulin secretion could be recovered by challenging stem cell-derived β-cells with intermediate metabolites from the TCA cycle and late (but not early) glycolysis; this resulted in robust, bi-phasic insulin release *in vitro* similar in magnitude to functionally mature human islets [41]. Another reason for the inability of hESC- and hiPSC-derived β-cells to respond to glucose may be that our and similar differentiation protocols yield polyhormonal cells (e.g. INS^+^ and GCG^+^ cells). Bruin et al [42] has provided comprehensive characterization of hESC-derived bihormonal cells: they show that these cells were responsive to KCl and arginine, but not glucose, in perifusion studies. An explanation for loss of glucose sensing in these cells could be lack of GLUT1 protein and reduced K_ATP_-channel activity. Indeed, they report that the expression of the SUR1 subunit of the K_ATP_-channel was ∼ 5-fold lower than KIR6.2). Combined, this suggests that an impaired ratio of SUR1:KIR6.2 may contribute to the observed K_ATP_-channel defects in hESC-derived islet endocrine cells, along with lack of GLUT1, which may underlie the absence of glucose-stimulated insulin secretion. Indeed, we observe expression of *GCG* in our hESC- and hiPS-derived b-cells, albeit at several fold lower levels than that of *INS*.

It is clear that the utilizing hiPSC as cellular systems to model disease mechanisms will become increasingly useful in biomedical research [43], including T2D research, where hiPSCs derived from humans could serve as versatile tools to understand mechanisms of human disease. There is strong evidence that the common *MNTR1B* variant (rs10830963) is associated with increased T2D risk [2, 16, 44–49], and perturbed β-cell function. Functional studies to determine the precise molecular mechanisms linking the *MTNR1B* variants to β-cell dysfunction are therefore required. Our approach to understand the molecular underpinnings of the pathogenetic effects of the *MTNR1B* risk SNP was to evaluate functional outcomes in isogenic hiPSCs and hESCs carrying risk and non-risk alleles. Our successful single-base gene editing approach by CRISPR/Cas9 enabled functional studies of the *MTNR1B* risk allele. However, differentiation protocols need to be improved and further developed to increase maturity and yield. Here, 3D- based protocols hold promise to generate islet cell-like clusters, resembling primary tissue (human islets of Langerhans), to determine direct causality of T2D risk alleles on β-cell function. Another limitation is that, at this point, the protocols for differentiation into β-cells are extremely labour-intensive: this restricts the number of cell preparations that can be generated for functional and expression studies, underpowering the experiments. This is a special concern when T2D risk alleles are examined and which presumably possess small effect sizes.

## Acknowledgements

We acknowledge the patients of the DIACT study, and the Lund Stem Cell Center Cell and the Gene Therapy Core funded by StemTherapy. Professor Johan Jakobsson for generation of fibroblasts and hiPSCs from patient skin biopsies. This work was supported by grants from the Swedish Research Council (2021-01116 to M.F.) and (2021- 01777 to H.M) the Novo Nordisk Foundation (NNF23OC0084475 - to M.F and NNF23OC0084463 - to H.M), the Hjelt Foundation (to M.F, S.K) and the Albert Påhlsson’s Foundation (to M.F.). The Crafoord Foundation (to S.K.). The European Commission (ERC- CoG_NASCENT – 681742 to P.W.F). The European Union’s Horizon 2020 research and innovation program (ISLET, no. 874839) (to AM and HS). Support was also received from the Lund University Diabetes Centre (LUDC) and the Swedish Foundation for Strategic Research (Dnr IRC15-0067 (to LUDC-IRC) and by the Swedish Research Council, Strategic Research Area Exodiab, Dnr 2009-1039.

## Author Contributions

Study design and conceptualization was by MF and HM, where PWF supported this process. Experiments were performed by TS, SK, JPMCMC, SH, FR, SG and AS. KYF and OE biopsied patients. AM and HS guided the development of the differentiation protocol. AR manages the DIACT study and performed GWAS and genotyping of patients in the study. MF wrote the manuscript together with TS, SK, SH and FR. All authors have read and approved the manuscript prior to submission.

## Figure and table legends

**Supplementary figure 1. Selected hiPSCs and hESCs are pluripotent.** (A-D) hiPSCs generated from fibroblasts and (E-G) a hESCs cell (HUES4) line show robust protein expression of pluripotency markers OCT4 (green) and TRA-1-81 (red). The scale bar represents 100 μm, and the images were captured at 10X magnification with a confocal microscope.

**Supplementary figure 2. MTNR1B protein is expressed in the pluripotent stem cell lines.** (A-D) hiPSCs and (C) hESCs show clear membrane protein expression of MTNR1B (green). The scale bar represents 100 μm, and the images were captured at 10X magnification with a confocal microscope.

**Supplementary figure 3. Tile scale images, immunocytochemistry and mRNA expression.** (A, B) hiPSCs and (C) hESCs successfully produced C-peptide (green) cells. Distribution of these cells can be observed over the entire surface area of a well. Images are captured at 10X magnification with a confocal microscope. Representative mages of C-peptide positive hiPSC derived β-cells (D), glucagon positive (E), and a merge (F). *GCG* gene expression in hiPSC derived (G) and hESCs derived (H) endocrine cells over the course of differentiation. The scale bar represents 100 μm, and the images were captured at 10X magnification with a confocal microscope.

**Supplementary figure 4. pLenti-HIP-GFP virus testing and titrations on clonal EndoC-βH1 line.** (A-F) Five days post lentivirus infection, cells showed robust expression of GFP. The scale bar represents 200 μm, and the images were captured at 10X magnification with a phase contrast microscope. (G-L) Percentage of GFP^+^ cells (bottom right corner) were assessed via flow cytometry and are presented as scatter plots. (G-I) From 1X to 4X dilution, more than 90% of cells expressed GFP under human insulin promoter (HIP).

**Supplementary figure 5. Sorting gating strategy for GFP^+^ (β-cells) from fully differentiated cells.** (A-C) hiPSCs and hESCs produced GFP^+^ β-cells at the end of differentiation after GFP-lentivirus infection; cells were selectively sorted from gate p4 (B’’’, C’’’) using FACS for downstream gene expression analysis. All panels show gating strategy in series to select cells based on (A-C) size-granularity (SSC-A vs FSC-A), and single cells (FSC-H vs FSC-A; SSC-H vs SSC-A). (A-A’’’) Panel A shows gating strategy on unstained (negative) control for GFP^+^ gate (p4) positioning.

## References

1. Ashcroft, F.M. and P. Rorsman, Diabetes mellitus and the beta cell: the last ten years. Cell, 2012. 148(6): p. 1160–71.

2. Lyssenko, V., et al., Common variant in MTNR1B associated with increased risk of type 2 diabetes and impaired early insulin secretion. Nat Genet, 2009. 41(1): p. 82–8.

3. Muoio, D.M. and C.B. Newgard, Mechanisms of disease:Molecular and metabolic mechanisms of insulin resistance and beta-cell failure in type 2 diabetes. Nat Rev Mol Cell Biol, 2008. 9(3): p. 193–205.

4. Suzuki, K., et al., Genetic drivers of heterogeneity in type 2 diabetes pathophysiology. Nature, 2024. 627(8003): p. 347–357.

5. Vujkovic, M., et al., Discovery of 318 new risk loci for type 2 diabetes and related vascular outcomes among 1.4 million participants in a multi-ancestry meta-analysis. Nat Genet, 2020. 52(7): p. 680–691.

6. Tuomi, T., et al., Increased Melatonin Signaling Is a Risk Factor for Type 2 Diabetes. Cell Metabolism, 2016. 23(6): p. 1067–1077.

7. Liu, J., et al., Melatonin Receptor 1B Genetic Variants on Susceptibility to Gestational Diabetes Mellitus: A Hospital-Based Case-Control Study in Wuhan, Central China. Diabetes Metab Syndr Obes, 2022. 15: p. 1207–1216.

8. Bouatia-Naji, N., et al., A variant near MTNR1B is associated with increased fasting plasma glucose levels and type 2 diabetes risk. Nature Genetics, 2009. 41(1): p. 89–94.

9. Prokopenko, I., et al., Variants in MTNR1B influence fasting glucose levels. Nature Genetics, 2009. 41(1): p. 77–81.

10. Vetter, C., et al., Night Shift Work, Genetic Risk, and Type 2 Diabetes in the UK Biobank. Diabetes Care, 2018. 41(4): p. 762–769.

11. Gaulton, K.J., et al., Genetic fine mapping and genomic annotation defines causal mechanisms at type 2 diabetes susceptibility loci. Nat Genet, 2015. 47(12): p. 1415–25.

12. Li, Y., H.V. Nguyen, and S.H. Tsang, Skin Biopsy and Patient-Specific Stem Cell Lines. Methods Mol Biol, 2016. 1353: p. 77–88.

13. Ameri, J., et al., Efficient Generation of Glucose-Responsive Beta Cells from Isolated GP2(+) Human Pancreatic Progenitors. Cell Rep, 2017. 19(1): p. 36–49.

14. Zufferey, R., et al., Multiply attenuated lentiviral vector achieves efficient gene delivery in vivo. Nat Biotechnol, 1997. 15(9): p. 871–5.

15. Cowan, C.A., et al., Derivation of embryonic stem-cell lines from human blastocysts. N Engl J Med, 2004. 350(13): p. 1353–6.

16. Tuomi, T., et al., Increased Melatonin Signaling Is a Risk Factor for Type 2 Diabetes. Cell Metab, 2016. 23(6): p. 1067–77.

17. Jiang, F. and J.A. Doudna, The structural biology of CRISPR-Cas systems. Curr Opin Struct Biol, 2015. 30: p. 100–111.

18. Nagorny, C.L., et al., Distribution of melatonin receptors in murine pancreatic islets. J Pineal Res, 2011. 50(4): p. 412–7.

19. Peschke, E., et al., Melatonin and type 2 diabetes - a possible link? J Pineal Res, 2007. 42(4): p. 350–8.

20. Zibolka, J., et al., Distribution and density of melatonin receptors in human main pancreatic islet cell types. J Pineal Res, 2018. 65(1): p. e12480.

21. Ramracheya, R.D., et al., Function and expression of melatonin receptors on human pancreatic islets. J Pineal Res, 2008. 44(3): p. 273–9.

22. Segerstolpe, A., et al., Single-Cell Transcriptome Profiling of Human Pancreatic Islets in Health and Type 2 Diabetes. Cell Metab, 2016. 24(4): p. 593–607.

23. Cardenas-Diaz, F.L., et al., Modeling Monogenic Diabetes using Human ESCs Reveals Developmental and Metabolic Deficiencies Caused by Mutations in HNF1A. Cell Stem Cell, 2019. 25(2): p. 273–289 e5.

24. Zhu, Z., et al., Genome Editing of Lineage Determinants in Human Pluripotent Stem Cells Reveals Mechanisms of Pancreatic Development and Diabetes. Cell Stem Cell, 2016. 18(6): p. 755–768.

25. Wang, X., et al., Point mutations in the PDX1 transactivation domain impair human beta-cell development and function. Mol Metab, 2019. 24: p. 80–97.

26. Velazco-Cruz, L., et al., Acquisition of Dynamic Function in Human Stem Cell-Derived beta Cells. Stem Cell Reports, 2019. 12(2): p. 351–365.

27. Rezania, A., et al., Reversal of diabetes with insulin-producing cells derived in vitro from human pluripotent stem cells. Nat Biotechnol, 2014. 32(11): p. 1121–33.

28. Pagliuca, F.W., et al., Generation of functional human pancreatic beta cells in vitro. Cell, 2014. 159(2): p. 428–39.

29. Peschke, E., Melatonin, endocrine pancreas and diabetes. J Pineal Res, 2008. 44(1): p. 26–40.

30. Peschke, E., et al., Receptor (MT(1)) mediated influence of melatonin on cAMP concentration and insulin secretion of rat insulinoma cells INS-1. J Pineal Res, 2002. 33(2): p. 63–71.

31. Kemp, D.M., M. Ubeda, and J.F. Habener, Identification and functional characterization of melatonin Mel 1a receptors in pancreatic beta cells: potential role in incretin-mediated cell function by sensitization of cAMP signaling. Mol Cell Endocrinol, 2002. 191(2): p. 157–66.

32. Picinato, M.C., et al., Melatonin inhibits insulin secretion and decreases PKA levels without interfering with glucose metabolism in rat pancreatic islets. J Pineal Res, 2002. 33(3): p. 156–60.

33. Peschke, E., A.G. Bach, and E. Muhlbauer, Parallel signaling pathways of melatonin in the pancreatic beta-cell. J Pineal Res, 2006. 40(2): p. 184–91.

34. Abdulwahab, D.A., et al., Melatonin protects the heart and pancreas by improving glucose homeostasis, oxidative stress, inflammation and apoptosis in T2DM-induced rats. Heliyon, 2021. 7(3): p. e06474.

35. Estaras, M., et al., Melatonin induces reactive oxygen species generation and changes in glutathione levels and reduces viability in human pancreatic stellate cells. J Physiol Biochem, 2019. 75(2): p. 185–197.

36. Lim, L.Y., et al., Generating pancreatic beta-like cells from human pluripotent stem cells. Methods Cell Biol, 2022. 170: p. 127–146.

37. Nair, G.G., et al., Recapitulating endocrine cell clustering in culture promotes maturation of human stem-cell-derived beta cells. Nat Cell Biol, 2019. 21(2): p. 263–274.

38. Hogrebe, N.J., et al., Targeting the cytoskeleton to direct pancreatic differentiation of human pluripotent stem cells. Nat Biotechnol, 2020. 38(4): p. 460–470.

39. Balboa, D., et al., Functional, metabolic and transcriptional maturation of human pancreatic islets derived from stem cells. Nat Biotechnol, 2022. 40(7): p. 1042–1055.

40. Barsby, T., et al., Differentiating functional human islet-like aggregates from pluripotent stem cells. STAR Protoc, 2022. 3(4): p. 101711.

41. Davis, J.C., et al., Glucose Response by Stem Cell-Derived beta Cells In Vitro Is Inhibited by a Bottleneck in Glycolysis. Cell Rep, 2020. 31(6): p. 107623.

42. Bruin, J.E., et al., Characterization of polyhormonal insulin-producing cells derived in vitro from human embryonic stem cells. Stem Cell Res, 2014. 12(1): p. 194–208.

43. Avior, Y., I. Sagi, and N. Benvenisty, Pluripotent stem cells in disease modelling and drug discovery. Nat Rev Mol Cell Biol, 2016. 17(3): p. 170–82.

44. Xia, Q., et al., Association between the melatonin receptor 1B gene polymorphism on the risk of type 2 diabetes, impaired glucose regulation: a meta-analysis. PLoS One, 2012. 7(11): p. e50107.

45. Kelliny, C., et al., Common genetic determinants of glucose homeostasis in healthy children: the European Youth Heart Study. Diabetes, 2009. 58(12): p. 2939–45.

46. Barker, A., et al., Association of genetic Loci with glucose levels in childhood and adolescence: a meta-analysis of over 6,000 children. Diabetes, 2011. 60(6): p. 1805–12.

47. Walford, G.A., et al., Common genetic variants differentially influence the transition from clinically defined states of fasting glucose metabolism. Diabetologia, 2012. 55(2):p. 331–9.

48. Langenberg, C., et al., Common genetic variation in the melatonin receptor 1B gene (MTNR1B) is associated with decreased early-phase insulin response. Diabetologia, 2009. 52(8): p. 1537–42.

49. Liao, S., et al., Association of genetic variants of melatonin receptor 1B with gestational plasma glucose level and risk of glucose intolerance in pregnant Chinese women. PLoS One, 2012. 7(7): p. e40113.

