## Supplementary figures and tables for "Modelling genetic risk of β-cell dysfunction in human induced pluripotent stem cells from patients carrying the *MTNR1B* risk variant"

**Supplementary Table 1. Antibodies used for immunocytochemistry**

| | Antibody | Catalogue number | Source | Dilution |
|---|---|---|---|---|
| 1 | Rabbit anti-OCT4 | PA5-27438 | Invitrogen | 1 in 500 |
| 2 | Mouse anti-TRA-1-81 | MA1-024 | Invitrogen | 1 in 500 |
| 3 | Rabbit anti-MTNR1B | PA5-102107 | Invitrogen | 1 in 500 |
| 4 | Rat anti-cPeptide | GN-1D4-s | DSHB* | 1 in 20 |
| 5 | Goat anti-PDX1 | ab47383 | Abcam | 1 in 5000 |
| 6 | Rat anti-NKX6.1 | F55A10 | DSHB | 1 in 500 |

- https://dshb.biology.uiowa.edu

**Supplementary Table 2. Taqman assays used for qRT-PCR**

| | TacMan Assays | Catalogue number | Source |
|---|---|---|---|
| 1 | *TBP* (housekeeping) | Hs00427620_m1 | ThermoFisher |
| 2 | *PPIA* (housekeeping) | Hs04194521_s1 | ThermoFisher |
| 3 | *MTNR1B* | Hs00173794_m1 | ThermoFisher |
| 4 | *INS* | Hs02741908_m1 | ThermoFisher |
| 5 | *PDX1* | Hs00236830_m1 | ThermoFisher |

**Supplementary figure 1.**

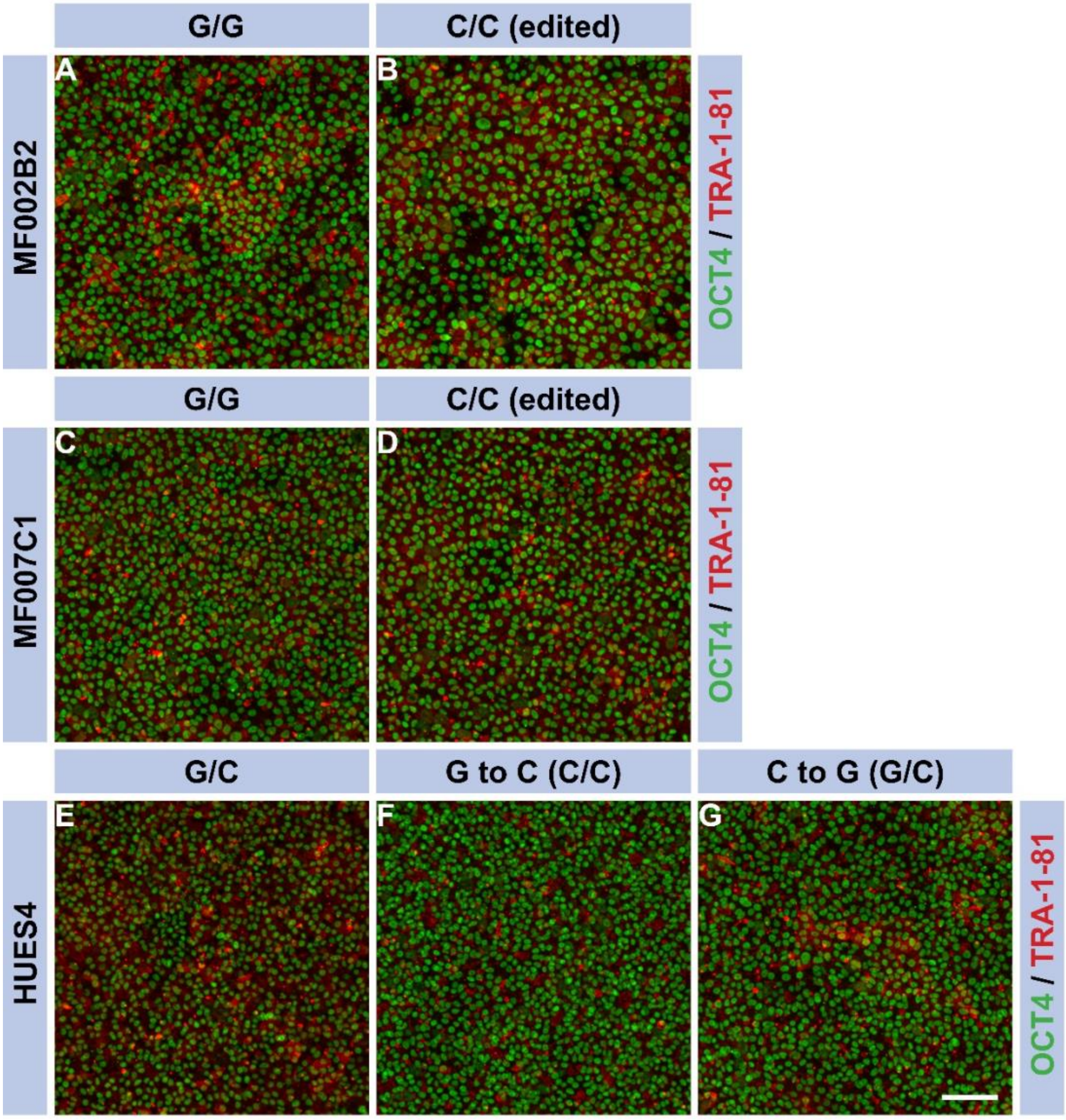

**Supplementary figure 2.**

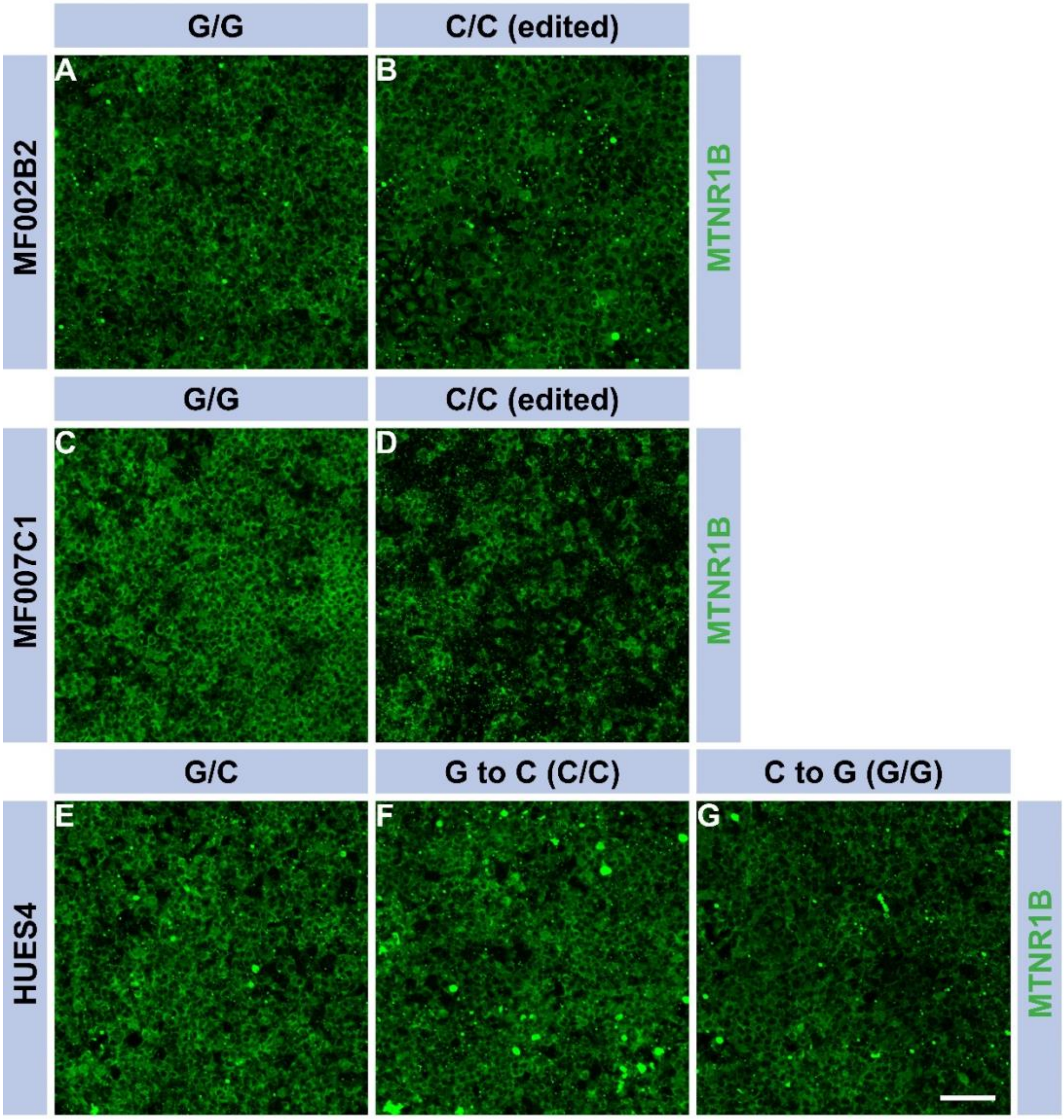

**Supplementary figure 3.**

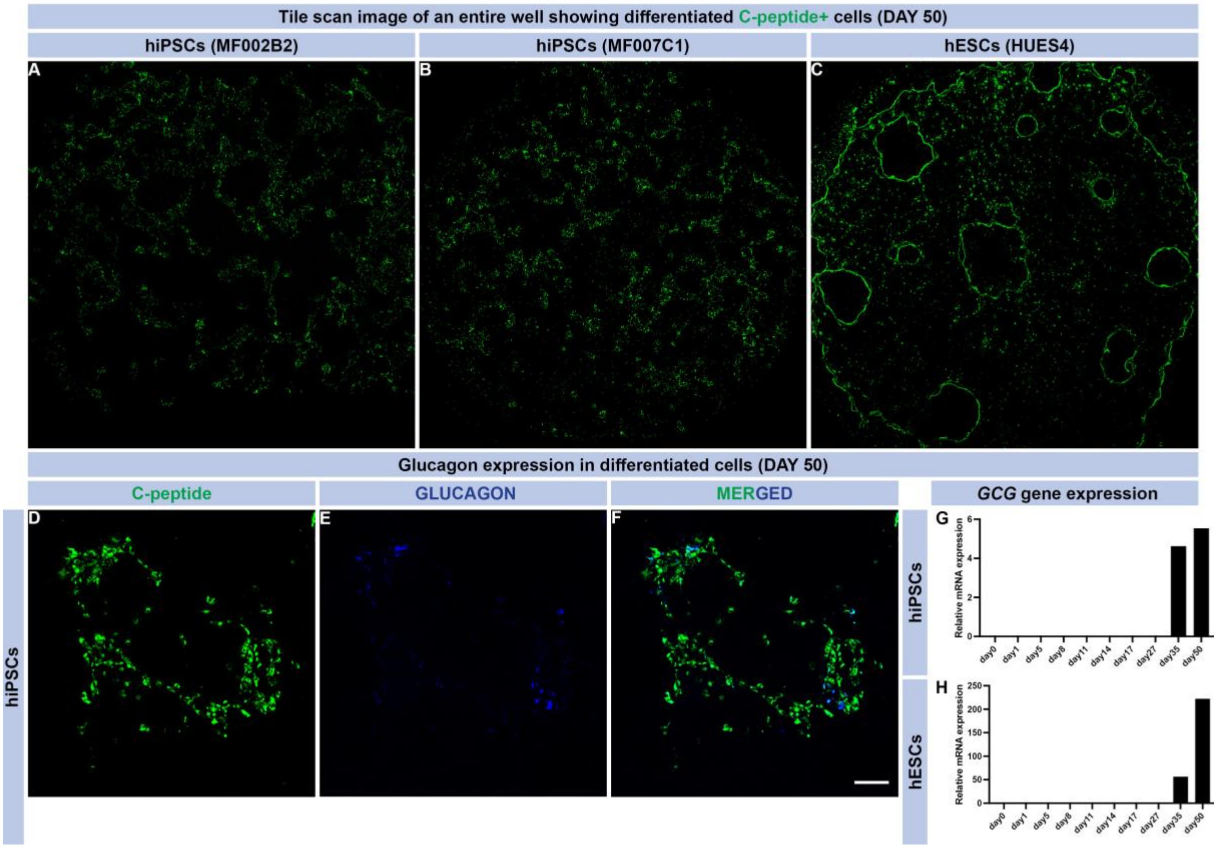

**Supplementary figure 4.**

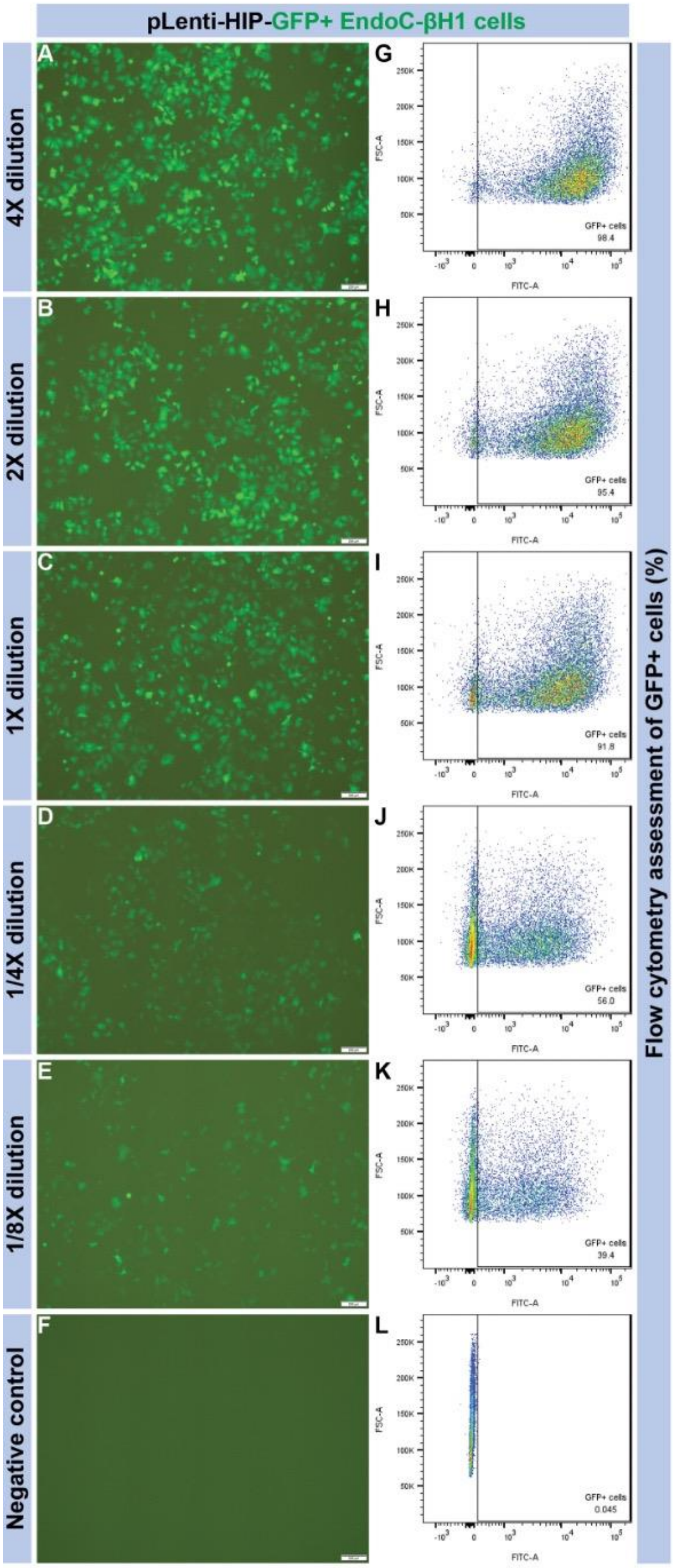

**Supplementary figure 5.**

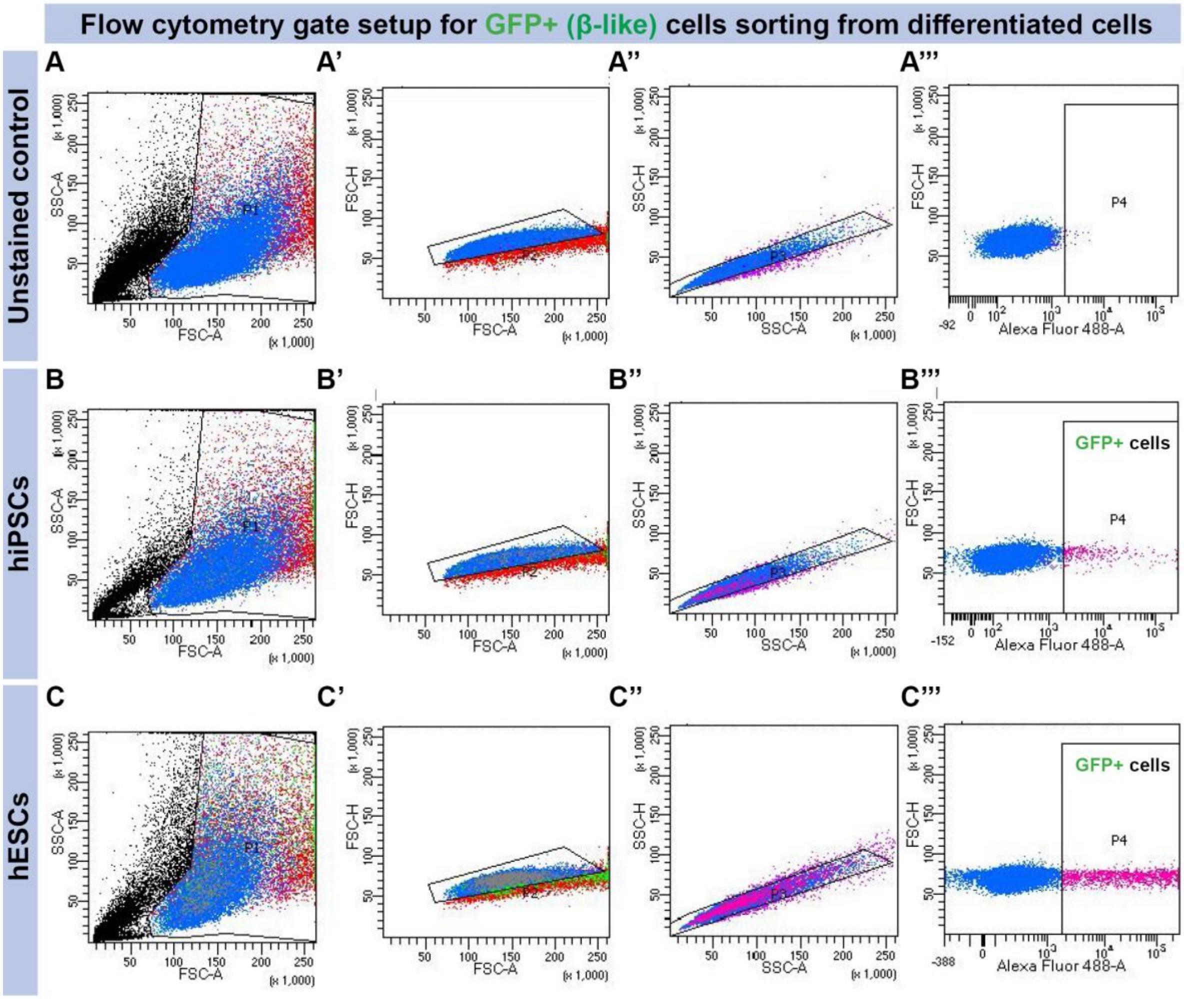
